# Glucose clearance and uptake is increased in the SOD1_G93A_ mouse model of amyotrophic lateral sclerosis through an insulin-independent mechanism

**DOI:** 10.1101/2020.08.02.233411

**Authors:** Tanya S. McDonald, Vinod Kumar, Jenny N. Fung, Trent M. Woodruff, John D. Lee

## Abstract

Metabolic disturbances are associated with the progression of the neurodegenerative disorder, amyotrophic lateral sclerosis (ALS), however the molecular events that drive energy imbalances in ALS are not completely understood. In this study we aimed to elucidate deficits in energy homeostasis in the SOD1_G93A_ mouse model of ALS. We identified that SOD1_G93A_ mice at mid-symptomatic disease stage have increased oxygen consumption and faster exogenous glucose uptake, despite presenting with normal insulin tolerance. Fasting glucose homeostasis was also disturbed, along with increased liver glycogen stores, despite elevated circulating glucagon, suggesting that glucagon signalling is impaired. Metabolic gene expression profiling of livers indicated that glucose cannot be utilised efficiently in SOD1_G93A_ mice. Overall, we demonstrate that glucose homeostasis and uptake are altered in SOD1_G93A_ mice, which is linked to an increase in insulin-independent glucose uptake and a disturbance in glucagon sensitivity, suggesting glucagon secretion and signalling could be potential therapeutic targets for ALS.

## Introduction

Amyotrophic lateral sclerosis (ALS) is a neurodegenerative disorder which is characterised by the progressive loss of motor neurons from both the cortex and spinal cord [1]. This loss of motor neurons leads to symptoms that are associated with ALS including muscle weakness, reduced motor control and the denervation and atrophy of skeletal muscle [1, 2]. ALS is a heterogeneous disease with the site of symptom onset varying amongst patients, and is known to have a complex etiology, with the exact cause of disease onset still unknown [3, 4]. However, disturbances in several pathways including dysfunctions in energy metabolism have been associated with the progression of ALS [5, 6].

Both sporadic and familial ALS patients are unable to maintain their body weight, with a loss of both fat and muscle mass shown early in the disease progression [7–9]. Rapid weight loss in patients is associated with worse disease outcomes, whilst conversely, a higher body mass index tends to increase survival rate [10, 11]. Evidence also suggests that insulin resistance plays a role in disease progression in both patients and animal models of ALS. For example, studies have shown an increased risk of ALS in people with diabetes metillus, and several animal models displayed reduced sensitivity to exogenous insulin [12–16]. However, there are also conflicting reports that have shown that premorbid type II diabetes delays the onset of symptoms in ALS patients, and some studies in both patients and mouse models have demonstrated glucose tolerance is unchanged [13, 17–20].

Studies examining metabolic dysfunction in ALS have mainly focussed on assessing how changes in glucose tolerance and insulin resistance affect metabolism in skeletal muscle and spinal cord [2, 13, 14, 17, 21]. In this study, we therefore aimed to elucidate the changes in whole body glucose homeostasis and metabolism using the SOD1_G93A_ mouse model of ALS, and identify some of the underlying disturbances in tissues such as the liver and pancreas, which are key drivers of regulating glucose homeostasis. Overall, we demonstrate that energy expenditure, and exogenous glucose uptake are increased through insulin-independent mechanism in SOD1 _G93A_ mice. Additionally, our findings indicate that glucagon signalling is impaired, leading to an accumulation of glycogen in the liver of SOD1_G93A_ mice. All of these perturbations in energy metabolism are likely to contribute to the dysregulation of glucose homeostasis in ALS.

## Results

### SOD1_G93A_ mice display loss of body weight and lean body mass with decreased activity and increased oxygen consumption at mid-symptomatic stage of disease

Using indirect calorimetry, we first aimed to delineate if the weight loss frequently observed in SOD1_G93A_ mice was due to a reduction in food intake, or increase in energy expenditure. At disease onset, SOD1_G93A_ mice showed no difference in body weight, but a 10% loss in lean body mass when compared to wild-type (WT) littermates. Furthermore, at mid-symptomatic stage, SOD1_G93A_ mice weighed significantly less than their WT littermates, with an 8 and 10% loss in total body weight and lean body mass respectively (Figure 1A and B). Despite this observed loss in body weight, total food intake was similar between WT and SOD1_G93A_ mice during the light and dark cycles at both disease stages (Figure 1C). In contrast to food intake, the average oxygen consumption in SOD1_G93A_ mice at mid-symptomatic stage was 121 and 113% higher than WTs during the light and dark cycles, respectively, while showing no differences at the onset stage (Figure 1D – 1G). This increase in oxygen consumption at mid-symptomatic stage was not due to an increase in locomotor activity, as we demonstrated a reduction of 18 and 25% in locomotor activity during the dark cycle at the onset and mid-symptomatic stages, respectively (Figure 1H – 1K). Interestingly, during the light phase, mid-symptomatic SOD1_G93A_ mice were 126% more active than their WT littermates. Despite these biphasic changes, total locomotor activity over a 24-hour period was not significantly different at either disease stages (Figure 1I and 1K). Although we saw differences in the lean body mass between WT and SOD1_G93A_ mice, no correlation was found between lean body mass and the average oxygen consumption over a 24-hour period (Figure 1L).

**Figure 1.**
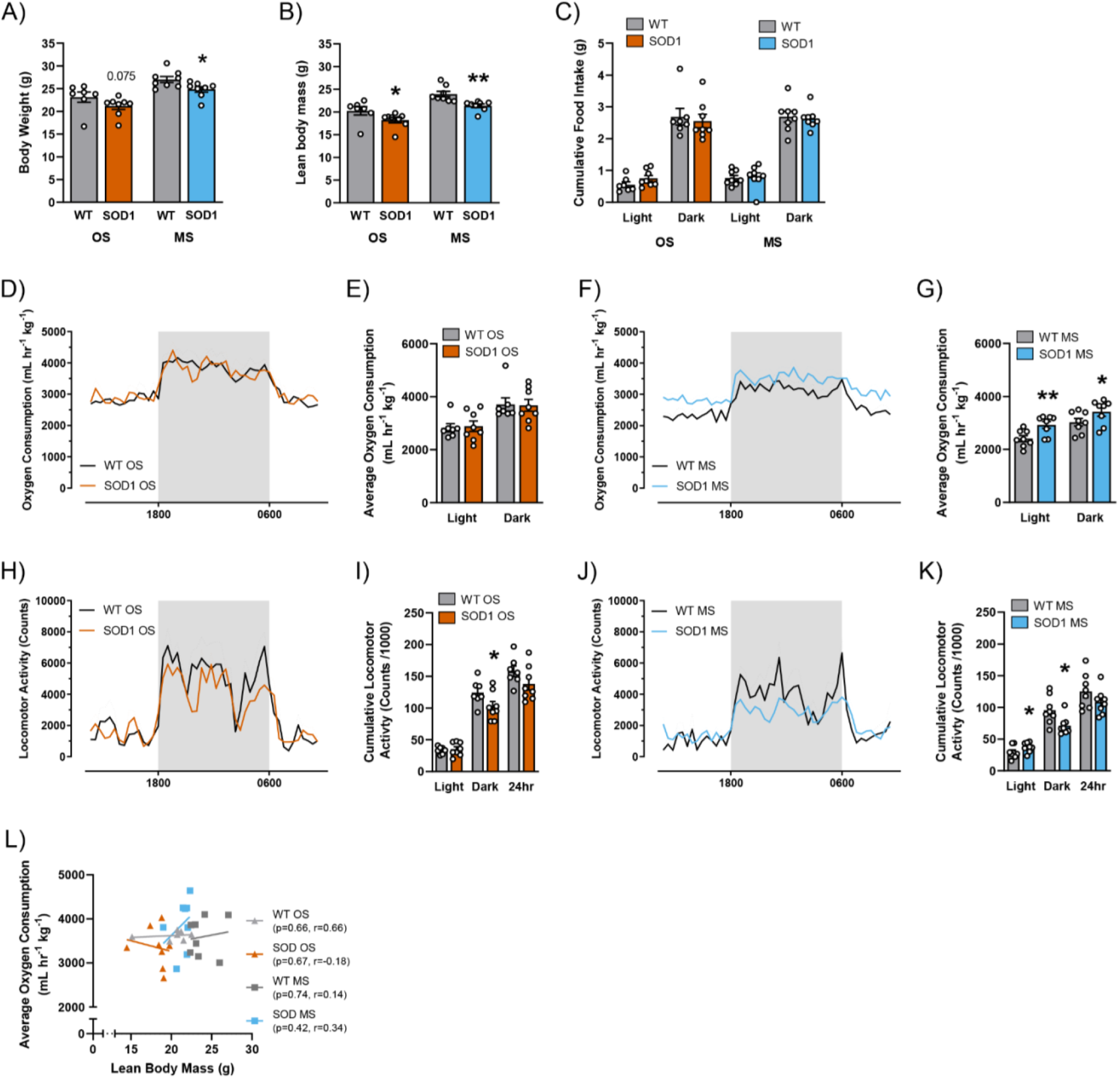
SOD1_G93A_ mice display loss of body weight and lean body mass associated with decreased activity and increased oxygen consumption. **(A, B)** Total body weight and lean mass of WT (grey bars) and SOD1_G93A_ (onset stage (OS) - orange bars and mid-symptomatic stage (MS) - blue bars) mice. **(C)** The cumulative food intake in both the light and dark phases, for WT (grey bars) and SOD1_G93A_ mice at OS (orange bars) and MS (blue bars). **(D-G)** Indirect calorimetry analysis was performed at both stages of disease using the Phenomaster TSE Metabolic Cage System. The 24-hour oxygen consumption profile in WT and SOD1_G93A_ mice at OS (**D**, WT – black lines, SOD1 _G93A_ – orange lines) and MS (**F**, WT – black lines, SOD1 _G93A_ – blue lines) is shown. From this profile the average oxygen consumption was calculated in both the light and dark cycle at OS (**E,** WT – grey bars, SOD1 _G93A_ – orange bars) and MS (**G,** WT – grey bars, SOD1 _G93A_ – blue bars). **(H-K)** The locomotor activity profile was assessed in WT and SOD1_G93A_ mice at OS (**H**, WT – black lines, SOD1 _G93A_ – orange lines) and MS (**J**, WT – black lines, SOD1 _G93A_ – blue lines). The cumulative locomotor activity in both SOD1_G93A_ and WT littermates were calculated for both the light and dark phases at OS (**I**, WT – grey bars, SOD1 _G93A_ – orange bars) and MS (**K**, WT black lines, SOD1 _G93A_ – blue lines). (**L**) Correlation analysis between lean body mass and the average oxygen consumption across the 24-hour period in WT (OS – light grey, MS – dark grey) and SOD1_G93A_ mice (OS – orange, MS – blue). No significance was found across genotype or stage of disease. All data presented as mean ± SEM; *n* = 8, for all measurements. All bar graphs analysed by two-tailed student *t*-test; * p < 0.05, ** p < 0.01.

### Exogenous glucose uptake is increased in SOD1_G93A_ mice at mid-symptomatic stage of disease

We next aimed to determine if glucose handling was altered in the SOD1_G93A_ mice by performing an intraperitoneal glucose tolerance test (ipGTT). At the onset of symptoms, SOD1_G93A_ mice and their WT littermates responded similarly to exogenous glucose (Figure 2A-E). However, at mid-symptomatic stage of disease, SOD1_G93A_ mice showed a faster rate of blood glucose clearance, with blood glucose levels being 33 to 50% lower in SOD1_G93A_ mice at 15, 30, 60- and 120-minutes post glucose injection when compared to WT mice (Figure 2F). This difference was further highlighted with a 46% drop in the area under the curve (AUC) of blood glucose levels (Figure 2G). As the glucose dose was calculated by weight, we wanted to confirm that the loss in body weight in SOD1_G93A_ mice was not responsible for the lower blood glucose concentrations throughout the ipGTT. Indeed, we showed that the reduction in the AUC of blood glucose levels did not correlate with the amount of glucose injected in these mice (Figure 2H and 2I). We also found that females had a similar ipGTT response to males, where SOD1_G93A_ mice at the mid-symptomatic stage had a 21% lower AUC of glucose concentration (Figure 2 – figure supplement 1A and 1B). To further validate this increase in blood glucose clearance, we intraperitoneally injected WT and SOD1_G93A_ mice with 2-deoxyglucose. Using LC-MS/MS, we found that 2-deoxyglucose uptake was 260 to 1040% higher in liver, brown and white adipose tissue of SOD1_G93A_ mice (Figure 2J). To elucidate the changes in glucose clearance of SOD1_G93A_ mice, we also measured plasma insulin and glucagon levels. Baseline plasma glucagon concentrations were 240% higher is SOD1_G93A_ mice compared to their WT littermates at the mid-symptomatic stage (Figure 2K). Although baseline insulin concentrations were unchanged, the plasma insulin response to exogenous glucose was significantly lower in SOD1_G93A_ mice, with a 44% reduction in AUC of insulin concentrations (Figure 2L and 2M). To confirm this reduction in insulin response, we also performed immunofluorescence of glucagon and insulin in the pancreas of WT and SOD1_G93A_ mice. We found a 22% reduction in the immunoreactive area of insulin-positive β-cells in the pancreas of SOD1_G93A_ mice when compared to WT mouse pancreas at mid-symptomatic stage (Figure 3A-3C), while there was no difference in the immunoreactive area of glucagon-positive α-cells at onset and mid-symptomatic stage of disease (Figure 3D and E).

**Figure 2.**
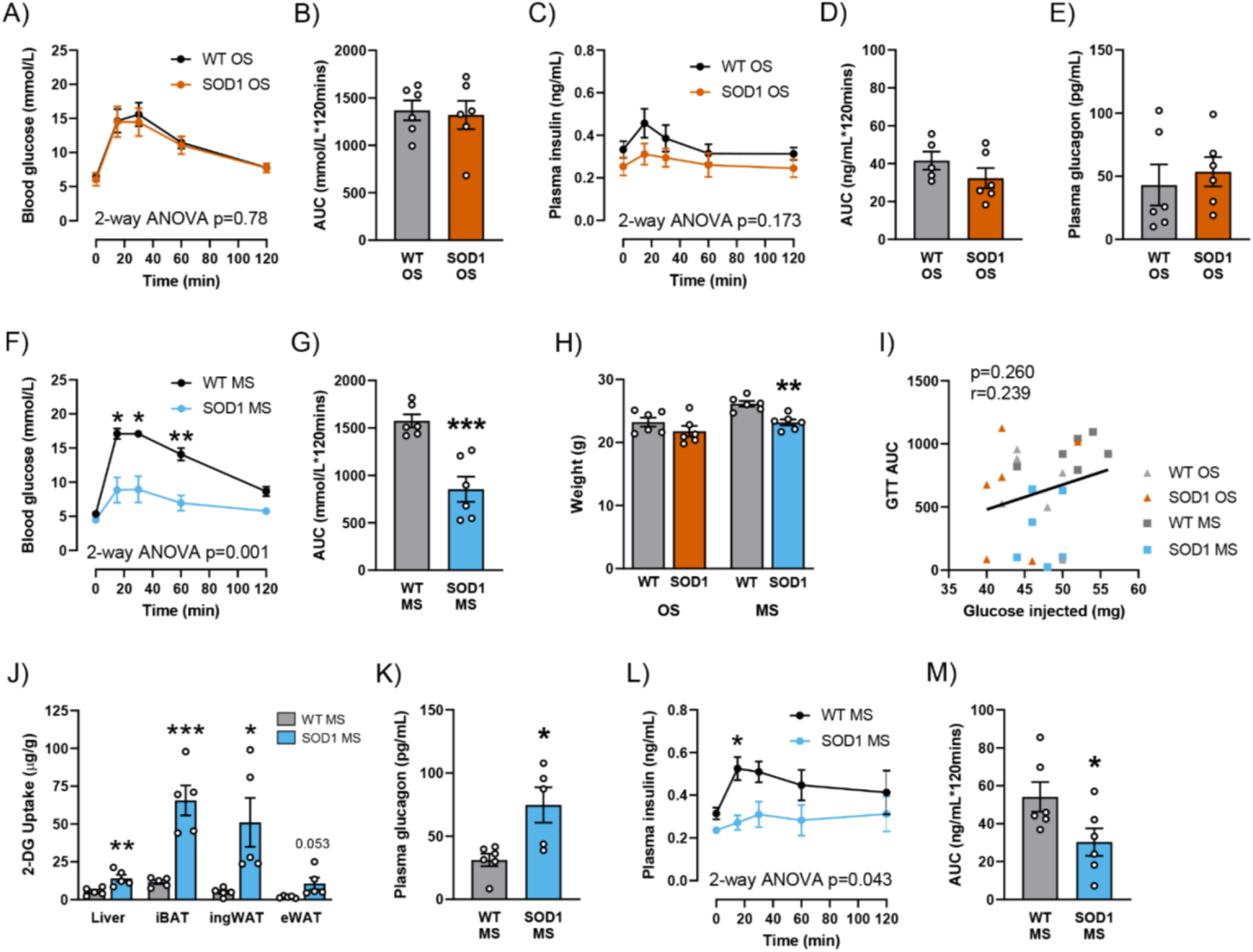
Glucose uptake is increased in SOD1_G93A_ mice, despite a reduction in plasma insulin levels at mid-symptomatic stage of disease. **(A)** Time course of blood glucose concentrations during a glucose tolerance test (ipGTT) following a 2 g/kg intraperitoneal injection of glucose at onset (OS) in WT (black dots) and SOD1_G93A_ (orange dots) mice. **(B)** The average area under the curve (AUC) calculated from the blood glucose time course for both WT (grey bars) and SOD1_G93A_ (orange bars) at OS. **(C)** Time course of insulin concentrations measured in plasma collected from tail bleeds throughout ipGTT at OS in WT (black dots) and SOD1_G93A_ (orange dots) mice. **(D)** The average AUC was calculated from the insulin time course as a measure of glucose-stimulated insulin release in WT (grey bars) and SOD1_G93A_ (orange bars) at OS. **(E)** Plasma glucagon levels measured from baseline tail bleeds from WT (grey bars) and SOD1_G93A_ (orange bars) at OS. **(F)** Time course of blood glucose concentrations during ipGTT in WT (black dots) and SOD1_G93A_ (blue dots) mice at mid-symptomatic (MS) stage of disease. **(G)** The average AUC calculated from the blood glucose time course for both WT (grey bars) and SOD1_G93A_ mice (blue bars) at MS. (**H**) Body weights of WT and SOD1_G93A_ mice used for ipGTT at OS and MS. **(I)** Pearson correlation analysis between the amount of glucose injected and the AUC of blood glucose concentrations. A single correlation analysis was performed on all WT and SOD1_G93A_ mice from both disease stages, represented by black line. (**J**) 2-deoxglucose uptake was measured in the liver, intrascapular brown adipose tissue (iBAT), inguinal white adipose tissue (ingWAT) and epididymal white adipose tissue (eWAT) following a 2 g/kg intraperitoneal injection of 2-deoxyglucose in fasted WT (grey bars) and SOD1_G93A_ mice (blue bars) at MS. **(K)** Plasma glucagon levels measured from baseline tail bleeds from WT (grey bars) and SOD1_G93A_ (blue bars) at MS. **(L)** Time course of insulin concentrations measured in plasma collected from tail bleeds throughout ipGTT in WT (black dots) and SOD1_G93A_ mice (blue dots) at MS. **(M)** The average AUC was calculated from the insulin time course as a measure of glucose-stimulated insulin release in WT (grey bars) and SOD1_G93A_ mice (blue bars) at MS. All data presented as mean ± SEM. *N* = 5-6, for all measurements. Two-way ANOVA results listed on time course in panels **A**, **C**, **F** and **L** is the overall significance between genotypes, Bonferroni *post-hoc* test was used to determine significant changes at specific time points. All bar graphs analysed by two-tailed student *t*-test. * p < 0.05, ** p < 0.01, *** p < 0.001.

**Figure 3.**
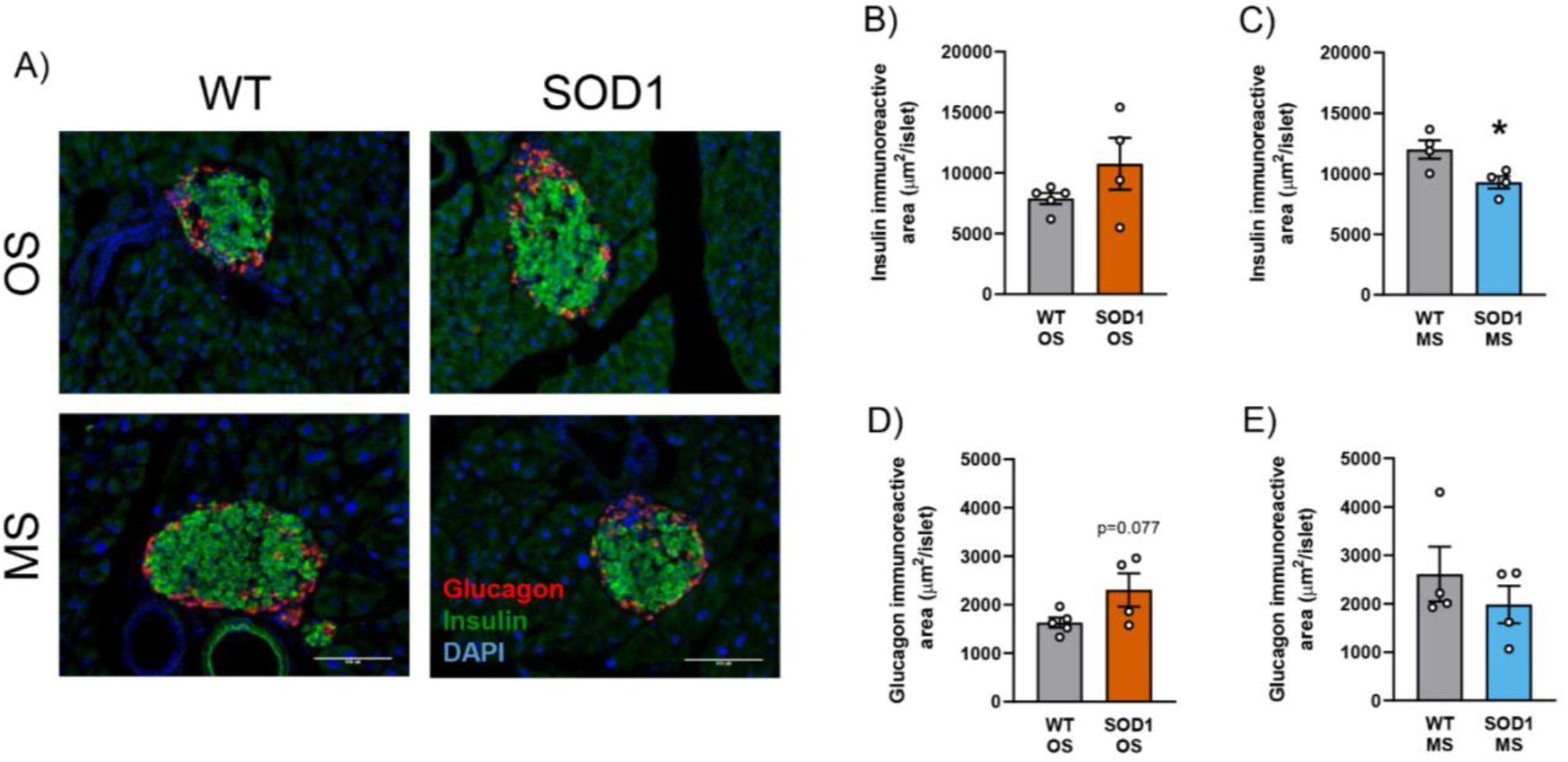
The area of β-cells in pancreatic islets are reduced in SOD1_G93A_ mice at mid-symptomatic stage of disease. **(A)** Representative images of glucagon-positive α-cells (red) and insulin-positive β-cells (green) in the pancreas of WT and SOD1_G93A_ mice at onset (OS) and mid-symptomatic (MS) stage of disease. **(B, C)** The average immunoreactive area of insulin-positive β-cells in the pancreatic islet cells at OS (**B**, WT – grey bars, SOD1 _G93A_ – orange bars) and MS (**C**, WT – grey bars, SOD1 _G93A_ – blue bars). **(D, E)** The average immunoreactive area of glucagon-positive α-cells in the pancreatic islet cells at OS (**D**, WT – grey bars, SOD1 _G93A_ – orange bars) and MS (**E**, WT – grey bars, SOD1 _G93A_ – blue bars). Scale bar = 100 μM. All data presented as mean ± SEM. *n* = 4-5. All bar graphs analysed by two-tailed student *t*-test. * p < 0.05.

### Insulin tolerance is unaffected in SOD1_G93A_ mice, despite a loss in fasting blood glucose concentrations

After a week of recovery from ipGTT, an intraperitoneal insulin tolerance test (ipITT) was performed on the same mice to determine insulin tolerance. At onset of disease, the response to exogenous insulin in SOD1_G93A_ mice was similar to that of their WT littermates (Figure 4A-E). Interestingly, SOD1_G93A_ mice had lower baseline blood glucose concentrations compared to WT mice following a 5-hour fast at mid-symptomatic stage (Figure 4F). Despite this difference in baseline blood glucose concentrations, both WT and SOD1_G93A_ mice responded similarly to insulin, as shown by the inverse AUC of blood glucose levels (Figure 4G). We also found changes in the blood hormone profile during the ipITT, with plasma glucagon concentrations trending higher at every timepoint measured, resulting in an overall 210% increase in AUC of glucagon concentrations for the SOD1_G93A_ mice (Figure 4H and 4I). Although a trend of decreased basal insulin concentrations was observed in the SOD1_G93A_ mice, no significance was observed (Figure 4J). Similarly, female mice at mid-symptomatic stage also did not show any difference in their ipITT response, but unlike males, no change in fasting blood glucose concentrations were found (Figure 4 – figure supplement 1A and 1B). Despite SOD1_G93A_ mice weighing less at both the onset and mid-symptomatic stage, the amount of insulin injected had no correlation with the inverse AUC of blood glucose levels (Figure 4K and 4L). In addition to insulin and glucagon levels, we also demonstrated that glycogen concentrations were 210 and 480% higher in the liver of SOD1_G93A_ mice at both onset and mid-symptomatic stages of disease, respectively (Figure 4M). Following an overnight fast, glycogen accumulation in the liver was still 400 to 500% higher in mid-symptomatic SOD1_G93A_ mice in comparison to their WT littermates (Figure 4M).

**Figure 4.**
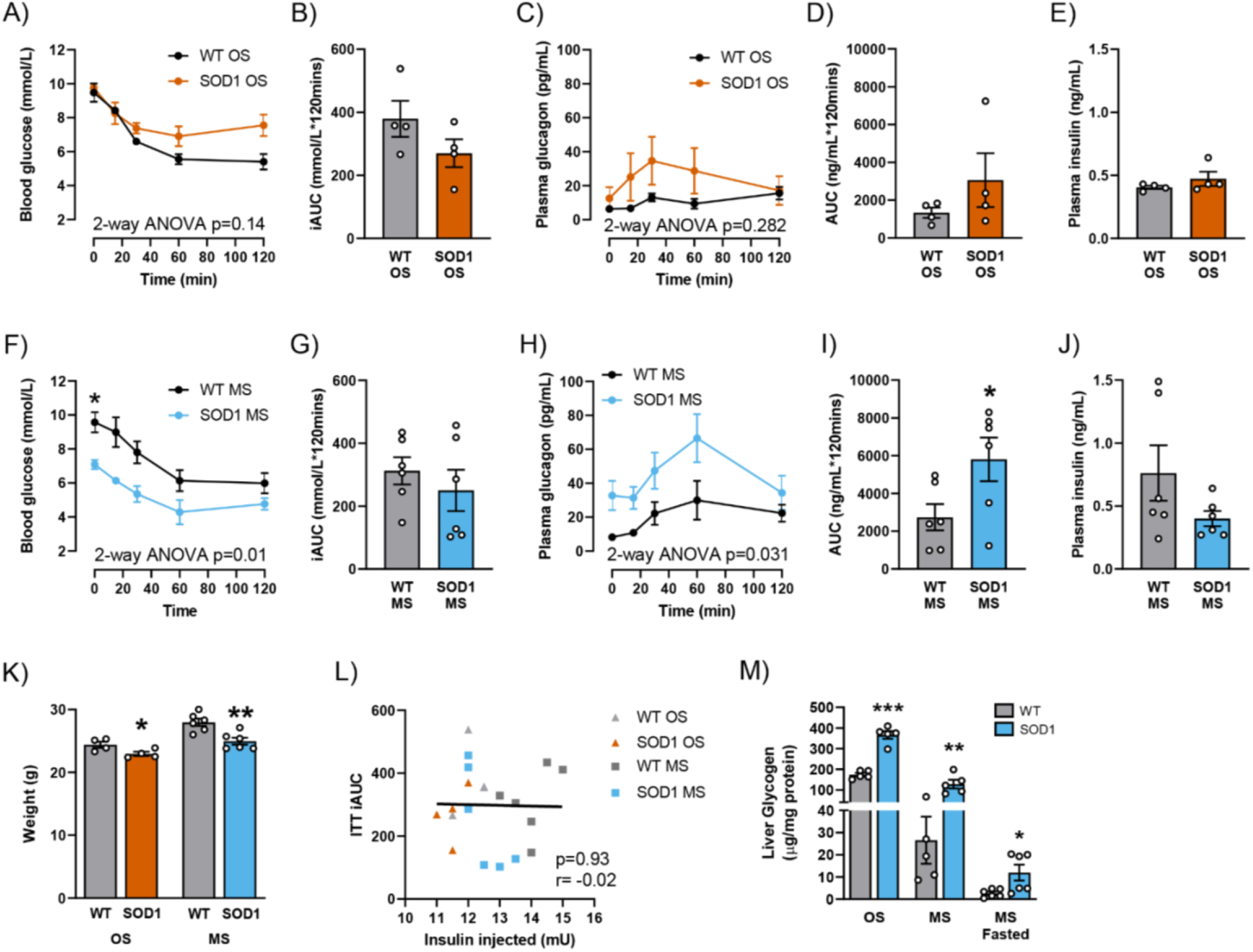
SOD1_G93A_ mice show a similar response to exogenous insulin, despite lower fasting blood glucose levels. **(A)** Time course of blood glucose concentrations during an insulin tolerance test (ipITT) at onset (OS) following a 0.75 IU/kg intraperitoneal injection of human insulin in WT (black dots) and SOD1 _G93A_ (orange dots). **(B)** The average inverse area under the curve (iAUC) was calculated from the blood glucose time course for both WT (grey bars) and SOD1_G93A_ (orange bars) at OS, to measure degree of insulin resistance. **(C)** Time course of plasma glucagon concentrations collected from tail bleeds throughout ipITT in both\WT (black dots) and SOD1_G93A_ mice (orange dots) at OS. **(D)** The average AUC was calculated from the glucagon time course for both WT (grey bars) and SOD1_G93A_ (orange bars) mice at OS. **(E)** Baseline plasma insulin levels were also measured from tail bleeds at OS in both WT (grey bars) and SOD1_G93A_ (orange bars) mice. **(F)** Time course of blood glucose concentrations during an ipITT at mid-symptomatic (MS) stage of disease following a 0.5 IU/kg intraperitoneal injection of human insulin in WT (black dots) and SOD1 _G93A_ mice (blue dots). **(G)** The average iAUC was calculated from the blood glucose time course for both WT (grey bars) and SOD1_G93A_ mice (blue bars) at MS. **(H)** Time course of plasma glucagon concentrations collected from tail bleeds in WT (black dots) and SOD1_G93A_ (blue dots) at MS. **(I)** The average AUC was calculated from the glucagon time course for both WT (grey bars) and SOD1_G93A_ mice (blue bars) at MS. **(J)** Baseline plasma insulin levels from tail bleeds at MS in both WT (grey bars) and SOD1_G93A_ mice (blue bars). **(K)** Body weights of WT (grey bars – both stages) and SOD1_G93A_ (OS – orange bars, MS – blue bars) mice used for ipITT. **(L)** Pearson correlation analysis between the amount of insulin injected (mU) and the iAUC of blood glucose concentrations during ipITT. Correlation analysis were performed on all WT (OS – light grey, MS – dark grey) and SOD1_G93A_ mice (OS – orange, MS – blue) from both disease stages together, represented by black line. **(M)** Glycogen concentrations were measured in liver homogenates at OS, MS, and overnight fasted MS (WT – grey bars and SOD1_G93A_ mice-light blue bars). All data presented as mean ± SEM. *n* = 4 at OS, and *n* = 6 at MS, for all measurements. Two-way ANOVA results listed on time course in panels **A**, **C**, **F** and **H** is the overall significance between genotypes, Bonferroni *post-hoc* test was used to determine significant changes at specific time points. All bar graphs analysed by two-tailed student *t*-test. * p < 0.05, ** p < 0.01, *** p < 0.001.

### Genes associated with glucose handling are altered in the liver of SOD1_G93A_ mice

To understand if this increase in glycogen was due to changes in the metabolic pathways in the liver, we conducted a metabolic PCR array consisting of 84 genes associated with glucose metabolism on livers from WT and SOD1_G93A_ mice at mid-symptomatic stage. The expression of twenty-four of these genes were significantly different between SOD1_G93A_ and WT mice, and are involved in one of four distinct pathways of glucose metabolism (Figure 5A). Firstly, two isoforms of phosphoglucose mutase, a reversible enzyme that catalyses glucose 6-phosphate to glucose 1-phosphate, were altered in SOD1_G93A_ mice, where *Pgm1* expression was significantly reduced by 0.7-fold while *Pgm3* expression was increased by 1.3-fold when compared to WT mice. In the glycogenesis pathway, UDP-glucose pyrophosphorylase 2 (*Ugp2,* 0.74-fold) and glycogen branching enzyme (*Gbe1*, 0.7-fold) expression were reduced in SOD1_G93A_ mice, whereas an isoform of glycogen synthase (*Gsk3a*, 1.6-fold), which inhibits glycogen synthase activity, was increased in SOD1_G93A_ mice. Overall, this indicates that the glycogenetic pathway is decreased at the mid-symptomatic stage of disease. In comparison, the expression of the liver isoform of glycogen phosphorylase (*Pygl*, 0.54-fold), which regulates glycogenolytic pathway was also reduced. However, the expression of three isoforms of phosphorylase kinase (*Phka1*, 1.9-fold; *Phkb*, 1.6-fold; *Phkg2*, 1.4-fold), that activate glycogen phosphorylase were higher in the liver of SOD1_G93A_ mice (Figure 5B).

**Figure 5.**
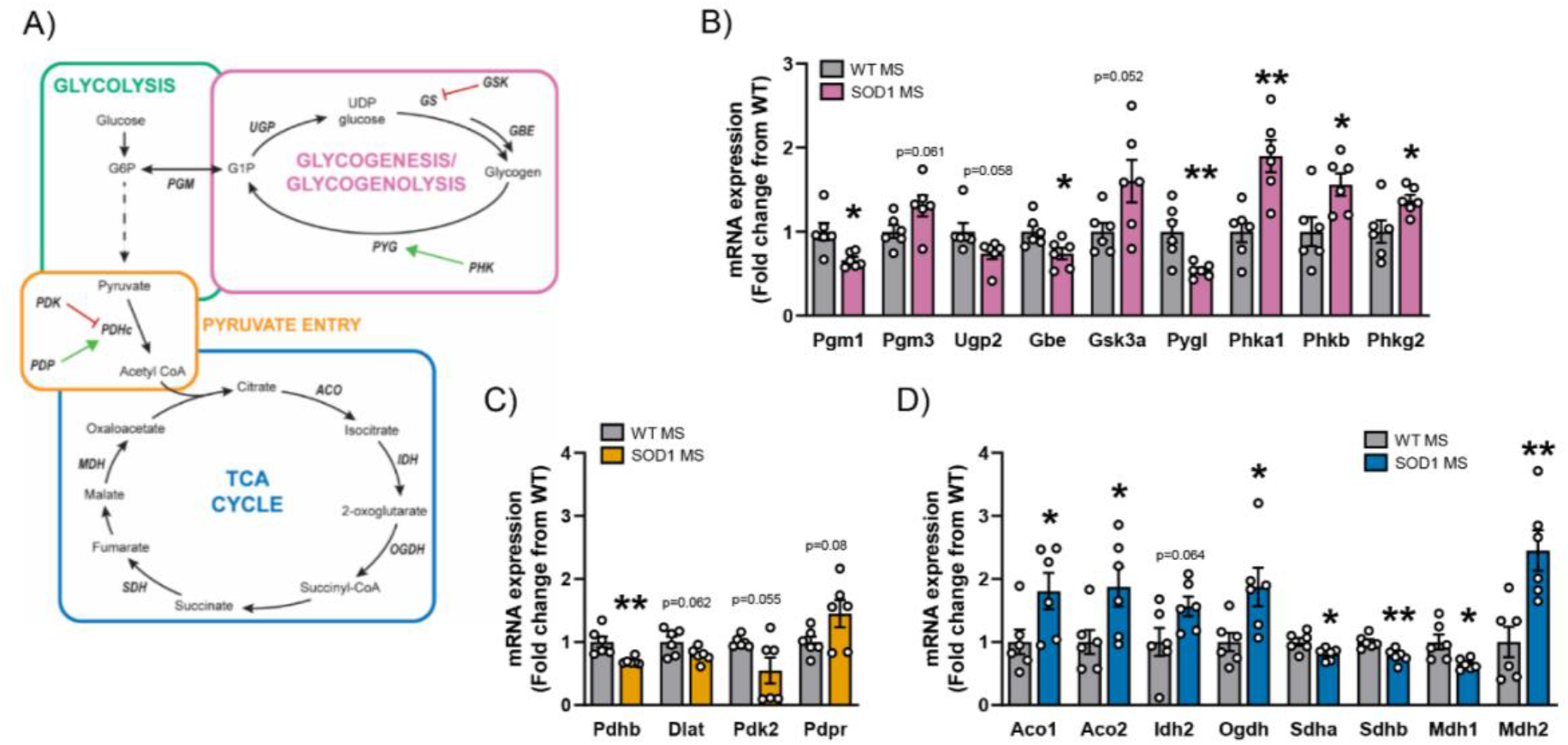
Glucose handling is altered in the liver of SOD1 _G93A_ mice at mid-symptomatic stage, potentially contributing to the increased liver glycogen stores found in both fed and fasted SOD1 _G93A_ mice. (**A**) Schematic of metabolic pathways that are changed in the liver of SOD1_G93A_ mice at mid-symptomatic (MS) stage of disease. **(B)** mRNA expression of genes involved in glycogenesis and glycogenolysis in the liver of WT and SOD1_G93A_ mice, which consists of phosphoglucose mutase (Pgm1 and Pgm3), UDP-glucose phosphorylase (Ugp2), glycogen branching enzyme (Gbe), and the liver isoform of glycogen phosphorylase (Pygl). As well as several genes that regulate activities of enzymes involved in these pathways including glycogen synthase kinase 3-alpha (Gsk3a) and three isoforms of phosphorylase kinase (Phka1, Phkb, and Phkg2). WTs represented by grey bars and SOD1_G93A_ mice by pink bars. **(C)** mRNA expression of genes involved in pyruvate entry into the TCA cycle in the liver of WT and SOD1_G93A_ mice at MS, including two of the enzymes that form the pyruvate dehydrogenase complex (PDHc), pyruvate dehydrogenase E1 subunit β (Pdhb) and dihydrolipoamide S-Acetyltransferase (Dlat), and genes that regulate PDHc activity, pyruvate dehydrogenase kinase (Pdk2) and pyruvate dehydrogenase phosphatase (Pdpr). WTs represented by grey bars and SOD1_G93A_ mice by yellow bars. **(D)** mRNA expression of key genes in the TCA cycle of WT and SOD1_G93A_ mice livers, including aconitase (Aco1, Aco2), isocitrate dehydrogenase (Idh2), oxoglutarate dehydrogenase (Ogdh), succinate dehydrogenase (Sdha, Sdhb) and malate dehydrogenase (Mdh1, Mdh2). WTs represented by grey bars and SOD1_G93A_ mice by dark blue bars. All data presented as mean ± SEM. *n* = 6, for all measurements. All bar graphs analysed by two-tailed student *t*-test. * p < 0.05, ** p < 0.01.

Genes associated with mitochondrial metabolism were also altered in the liver of SOD1_G93A_ mice at the mid-symptomatic stage of disease. Two of the three enzymes that comprise the pyruvate dehydrogenase complex, pyruvate dehydrogenase E1 subunit β (*Pdhb*, 0.7-fold) and dihydrolipoamide S-acetyltransferase (*Dlat*, 0.8-fold), were lower in the SOD1_G93A_ mice when compared to WT mice. Conversely, the expression of an isoform of pyruvate dehydrogenase kinase (*Pdk2*, 0.55-fold) which inhibits pyruvate dehydrogenase was reduced in SOD1_G93A_ mice, while pyruvate dehydrogenase phosphatase (*Pdpr*, 1.5-fold) that activates pyruvate dehydrogenase is increased in SOD1_G93A_ mice (Figure 5C). In contrast to the other pathways, the genes associated with TCA cycle activity were elevated in the liver of SOD1_G93A_ mice. Most of the enzymes in TCA cycle were upregulated in the liver of SOD1_G93A_ mice, including aconitase (*Aco1*, 1.8-fold; *Aco2*, 1.9-fold), isocitrate dehydrogenase (*Idh2*, 1.6-fold), oxoglurate dehydrogenase (*Ogdh*, 1.9-fold), and an isoform of malate dehydrogenase (*Mdh2*, 2.4-fold). Only the two subunits of succinate dehydrogenase (*Sdha*, 0.8-fold; *Sdhb*, 0.78-fold) and another isoform of malate dehydrogenase (*Mdh1*, 0.65-fold) were reduced in SOD1_G93A_ mice (Figure 5D). Taken together, this suggests that glycogenesis/glycogenolysis and pyruvate entry in the TCA cycle are decreased, while the TCA cycle is increased in the liver of SOD1_G93A_ mice. In addition, we demonstrated that 4 main clusters with similar gene expression profiles were evident using 34 genes that showed changes in expression between WT and SOD1_G93A_ mouse liver tissues with a p-value less than 0.1 (Figure 5-supplement figure 1A). These genes were analysed in the GEN2FUNC module of the Functional Mapping and Annotation of Genome-Wide Association (FUMA) software. The top 3 significant hallmark pathways (adjusted p-value < 10^−10^) enriched for gene expression included “Oxidative Phosphorylation (OXPHOS)”, “Hypoxia” and “Glycolysis” (Figure 5-supplement figure 1B).

## Discussion

The major finding of the current study is that there is a shift in energy balance in SOD1_G93A_ mice, and this shift in energy expenditure contributes to an increase in glucose clearance. This change in glucose clearance could be due to the increase in glucose uptake in the liver and adipose tissue in an insulin-independent manner. We concluded that these changes are insulin-independent as we found no difference in glucose disposal in response to exogenous insulin. Furthermore, we also demonstrated that SOD1_G93A_ mice have a reduction in insulin-positive β-cell area, and an impairment in the release of insulin in response to exogenous glucose. SOD1_G93A_ mice also showed an accumulation of glycogen in the liver, despite an increase in circulating glucagon concentrations and gene expression data, suggesting a decrease in both glycogen synthesis and breakdown. This indicates that glucagon signalling may be impaired in the liver of SOD1_G93A_ mice. Finally, the gene expression profile of several metabolic enzymes suggested that the liver switches from using glucose to fatty acids as an energy source, which has previously been found in skeletal muscle and CNS tissues in ALS.

Consistent with several other studies, we found that SOD1_G93A_ mice had reduced body weight, when compared to their WT littermates. At the mid-symptomatic stage of disease, oxygen consumption is increased in SOD1_G93A_ mice, independent of changes in both food intake and activity. This indicates that during disease progression, a shift occurs whereby energy expenditure increases in SOD1_G93A_ mice. Indeed studies have shown that a high fat diet, that supplies ALS mice with higher calories, increases survival, whilst calorie restriction accelerates disease progression [22]. Overall, this indicates that this change in energy balance at the later phase of the disease contributes to disease progression and survival.

To establish how alterations in whole-body metabolism contribute to the changes in energy expenditure during disease progression, we next assessed the response to both exogenous glucose and insulin using ipGTT and ipITT. We found that blood glucose clearance was more efficient under fasting conditions in SOD1_G93A_ mice at the mid-symptomatic stage. This was further validated in a separate cohort of mice, where 2-deoxyglucose uptake was higher in both liver and adipose tissues of SOD1_G93A_ mice. These results support previous findings in SOD1_G86R_ transgenic male ALS mice, in which an increased rate of glucose clearance and higher 2-deoxyglucose uptake was observed in white adipose tissue [22]. However, the results of recent studies using SOD1_G93A_ mice contradict our findings. One study found no difference in glucose clearance across all stages of disease, whereas, another two studies found reduced glucose clearance [13, 17, 21]. This discrepancy in glucose clearance echoes what is found in ALS patients, with numerous studies showing both reduced and unchanged glucose tolerance [18, 23, 24]. These inconsistencies in glucose homeostasis are also observed on the tissue level, where glucose uptake is increased in denervated forearm and skeletal muscle of ALS patients [25, 26]. In contrast, glucose metabolism was reduced in the cerebral cortex, and spinal cords of both patients and animal models [12, 13, 27–29]. This disparity in how glucose uptake and metabolism is altered in individual tissues may underlie the variety of responses we observe in whole-body glucose clearance. Overall, changes in glucose tolerance are complex in ALS, and still require further refined investigation.

In contrast to the ipGTT, we found the response to the ipITT was similar between SOD1_G93A_ and WT mice, indicating that insulin sensitivity is unaltered in these mice. This corroborates one study in ALS patients which found insulin sensitivity was unchanged during a euglycemic insulin clamp [30]. In an alternate study, it was demonstrated that the glucose infusion rate required to maintain glycemia was significantly lower in ALS patients compared to healthy controls, and thus these patients were considered insulin resistant [12]. Furthermore, in isolated fast twitch skeletal muscles from both SOD1_G93A_ and TDP43_A315T_ mice, insulin-stimulated glucose uptake was decreased [13, 14]. However, at the later stages of disease, insulin-independent glucose uptake was increased in the fast twitch muscle of SOD1_G93A_ mice, suggesting that insulin resistant muscle fibres may be compensating by upregulating pathways involved in insulin-independent glucose uptake [13]. A limitation of our study is that we cannot determine which tissues are specifically transporting glucose in response to exogenous insulin. However, as we see no change in glucose response to insulin, but an increase in glucose clearance in response to exogenous glucose, it suggests that although insulin signalling is intact, insulin-independent glucose uptake is elevated in the SOD1_G93A_ mice at the mid-symptomatic stage of disease.

To find the potential causes of the change in glucose disposal at the mid-symptomatic stage we analysed the concentrations of the glucoregulatory hormones, insulin and glucagon. At the mid-symptomatic stage of disease there was a reduced spike in plasma insulin concentrations in response to exogenous glucose. This loss in insulin may be due to the rapid uptake of glucose. Specifically, the change in blood glucose concentrations was not high enough or prolonged enough to stimulate insulin release from β-cells. However, in other studies it has been shown that ALS patients had a reduced insulin response during an oral GTT, with these studies concluding that there is an impaired synthesis or release of insulin due to pancreatic islet cell damage [24]. This is further supported by our demonstration of reduced insulin-positive β-cells in the pancreas of SOD1_G93A_ mice, suggesting that insulin synthesis and release is lower in SOD1_G93A_ mice. Together this further indicates that insulin-independent glucose uptake is upregulated in mid-symptomatic SOD1_G93A_ mice.

In comparison to insulin, glucagon concentrations were elevated in fasted SOD1_G93A_ mice and in response to the falling blood glucose during the ipITT. In ALS patients it was found that an increase in disease severity correlated with higher circulating glucagon concentrations [31]. Despite increased circulating glucagon, we still observed reduced fasting glucose concentrations in the ipITT. This is similar to ALS patients where they observed an increase in both fasting and postprandial glucagon concentrations, coupled with reduced fasting glucose [32]. Significantly decreased blood glucose concentrations in the presence of higher circulating glucagon suggests that both ALS patients and SOD1_G93A_ mice are glucagon insensitive. This could be caused by either a reduction in the expression of the glucagon receptor, or a loss in glucagon signalling in the liver. The primary role of glucagon in the liver is to stimulate glycogenolysis, the breakdown of glycogen to glucose, and the release of glucose into the circulation [33]. Interestingly, we detected increased glycogen accumulation in the liver even prior to a significant elevation in plasma glucagon concentrations. Shorter fasting periods are known to rely heavily on the glycogenolytic pathway to maintain blood glucose concentrations, whereas a prolonged fast of over 12 hours relies more on gluconeogenesis as glycogen stores become depleted [34]. This would help explain why a difference in fasting glucose concentrations in SOD1_G93A_ mice were observed after a 5-hour fast, but not an overnight fast. In this study we were unable to determine the cause of increased circulating glucagon, but it would be of interest to follow this further to understand how glucagon synthesis, secretion and signalling is altered in ALS.

The liver is crucial for maintaining glucose homeostasis, as it is responsible for both producing glucose during fasting, and storing glucose following a meal. Since we observed marked changes in glucose homeostasis in SOD1_G93A_ mice we investigated the changes in the expression of genes involved in metabolic pathways in the liver. At the mid-symptomatic stage of disease, the gene expression profile indicated that both glycogenesis and glycogenolysis is reduced in the liver. Glycogen accumulation has previously been shown in the liver as well as skeletal muscle and spinal cord throughout the course of disease in SOD1_G93A_ mice [35]. As mentioned above, this accumulation of glycogen may be due to a lack of glucagon signalling in the liver of SOD1_G93A_ mice. Along with a change in glycogen handling, entry of glucose into the TCA cycle appears to be reduced in SOD1_G93A_ mice, as the expression levels of two of the three enzymes that form the pyruvate dehydrogenase complex, which regulates pyruvate entry into the TCA cycle, are lower in the liver. Furthermore, the expression of several enzymes involved in the TCA cycle were upregulated in SOD1_G93A_ mice. Together this supports findings in other affected tissues that show a switch from using glucose to lipid as the primary fuel source for the TCA cycle [21, 36, 37]. Although the exact trigger that leads to this switch is unknown, it has been proposed that an increase in fatty acid metabolism occurs to compensate for the inability for tissues to use glucose and glycogen as energetic substrates. Although it is a beneficial short-term compensatory mechanism, chronic reliance on the metabolism of fatty acids via β-oxidation can lead to the accumulation of toxic by-products, in particular, reactive oxygen species (ROS). An increased production in ROS has been shown to proceed muscle denervation in the tibialis anterior of SOD1_G86R_ mice. Although it has yet to be investigated if elevated ROS production occurs in other peripheral tissues in ALS, it is clear that excessive ROS production can be deleterious in ALS [37]. Overall, changes in energy substrate usage in the liver and a change in glucoregulatory hormone secretion and signalling may underlie the impairments observed in glucose homeostasis in SOD1_G93A_ mice.

## Conclusion

Our findings suggest that glucose homeostasis and whole-body metabolism are relatively unchanged at the onset of symptoms in SOD1_G93A_ mice. However, throughout disease progression we identified an increase in glucose uptake in SOD1_G93A_ mice, likely occurring through insulin-independent mechanisms. This increased glucose uptake however was stored as glycogen in tissues such as the liver, rather than used as an energy source. Our results also indicate that SOD1_G93A_ mice may develop glucagon resistance, which has been reported previously in ALS. Overall our study documents that SOD1_G93A_ mice develop imbalances in glucose homeostasis, and that the secretion and signalling pathways of both glucagon and insulin may be potential therapeutic targets for ALS.

## Materials and Methods

### Ethical Statement

All experimental procedures were approved by the University of Queensland Animal Ethics Committee and complied with the policies and regulations regarding animal experimentation. They were conducted in accordance with the Queensland Government Animal Research Act 2001, associated Animal Care and Protection Regulations (2002 and 2008), and the Australian Code of Practice for the Care and Use of Animals for Scientific Purposes, 8th Edition. Animal studies are reported in compliance with the ARRIVE guidelines

### Animals

Transgenic SOD1_G93A_ mice (B6-Cg-Tg (SOD1-G93A) 1Gur/J) were obtained from the Jackson laboratory (Bar Harbor, ME, USA) and were bred on C57BL/6J background to produce SOD1_G93A_ and wild-type (WT) mice. SOD1_G93A_ mice at two predetermined stages of disease were used for this study; onset (OS, 70 days postnatal), the first sign of motor deficits as a significant decline in hind-limb grip strength and mid-symptomatic (MS, 130 days postnatal), where hind-limbs are markedly weaker and a tremor is observed when mouse is suspended by the tail [38]. The breeding colony was maintained at the University of Queensland Biological Resources Animal Facilities. Mice were housed under 12-hour light/dark cycle with free access to food and water unless otherwise stated.

### Assessment of body weight, food intake, activity and metabolism

Mice were singly housed and acclimated to metabolic chamber (TSE systems, Bad Homburg Germany) for 1 week prior to recordings. Animals were continuously recorded for 48 hours, with food intake, locomotor activity (in X and Y axes) and gas exchange (O_2_ and CO_2_) measured every 30 minutes for using the Phenomaster open-circuit indirect calorimetry system. Our analysis was performed on the final 24 hours of recording to allow mice to acclimatise to the Phenomaster system for the first 24 hours. Mice were maintained on 12 hours light/ dark cycle (on at 0600 h and off at 1800 h) with free access to food and water. Lean body mass of WT and SOD1_G93A_ mice were measured using Bruker Minispec LF50 Body Composition Analyser (7.5 MHz, Bruker Optics, Inc). Specifically, animals were placed in a clear plastic cylinder and kept immobile by inserting a tight-fitting plunger into the cylinder. The tube was inserted into the sample chamber of the instrument and scanned for approximately 2 minutes.

### Glucose/Insulin Tolerance Tests

For glucose/insulin tolerance tests, SOD1_G93A_ mice were housed with WT littermate on purachip bedding. To reduce stress, mice were handled daily for a week prior to tests. Intraperitoneal glucose tolerance tests (ipGTT) were performed after fasting mice overnight for 16 hours. Basal blood glucose levels were measured from a tail bleed using a Freestyle Optium Neo glucometer. Blood was then sampled from tail bleeds 15, 30, 60 and 120 minutes after intraperitoneal injection of glucose (2 g/kg) in 0.9% saline. At each time point approximately 50 μL of tail blood was collected in lithium-heparin coated capillary tubes. The capillary tubes were centrifuged at 2000 × *g* for 10 minutes at 4°C, and plasma collected and stored at −80°C until used.

A week after the ipGTT, intraperitoneal insulin tolerance tests (ipITT) were performed following a similar protocol. Mice were fasted for 6 hours in the morning, with blood collections and blood glucose measurements conducted at the same time points as above, following an intraperitoneal injection of 0.75 IU/kg human insulin at onset, and 0.5 IU/kg at mid-symptomatic stage. Initially, we tried a dose of 0.5 IU/kg of insulin at onset stage, however, at this age mice unexpectedly responded with an initial spike in blood glucose levels. This was independent of both sex and genotype (*results not shown*). Hence, using a separate cohort of mice, we tested a higher dose of 0.75 IU/kg, and at this dose we observed the expected decline in blood glucose concentrations suggesting that mice at this age may be less sensitive to human insulin under the specific conditions of the ipITT. The higher dose was used for onset stage to analyse changes in blood hormone concentrations.

### Enzyme-Linked Immunosorbent Assay (ELISA)

Plasma insulin and glucagon concentrations were measured using an Ultra-Sensitive Mouse Insulin ELISA Kit and Mouse Glucagon ELISA Kit (Crystal Chem, Illinois USA), respectively. Protocols were followed as per Manufacturer’s instructions. The endpoint colorimetric assays were performed using a Tecan Spark multimode microplate reader (Tecan, Mannedorf Switzerland). Area under the curve (AUC) was calculated individually for each mouse using the trapezoid rule. For ipITT blood concentrations over time, an inverse AUC (iAUC) was performed using basal blood glucose concentrations as baseline.

### 2-deoxyglucose uptake analysis

Mice were fasted overnight for 16 hours to replicate condition of ipGTT. Liver, intrascapular brown adipose tissue, inguinal white adipose tissue and epididymal white adipose tissue were collected 30 minutes after an intraperitoneal injection of 2-deoxyglucose (2 g/kg) in 0.9% saline. All tissues were store at −80°C until used.

Tissues were homogenised in a volume of milliQ water equivalent to tissue weight, spiked with [U-^13^C]-glucose as an internal standard. Polar metabolites were extracted using a Bligh-Dyer water/methanol/chloroform extraction procedure previously optimised [39]. The upper polar phase was freeze dried using Labconco centrivap cold trap (Labconco, Missouri USA), and then analytes were derivatised using O - benzylhydroxylamine hydrochloride (O-BHA) and N-(3-dimethylaminopropyl) - N′ - ethylcarbodiimide hydrochloride (EDC) under the conditions previously optimised [40]. Following derivatisation, the lower organic layer was evaporated using a nitrogen dryer at 40°C, and reconstituted in 100 μl of 50% methanol. Standards were prepared similarly in matrix free milliQ water.

The LC-MS/MS system consisted of an API 3200 (AB SCIEX) triple quadrupole LC-MS/MS mass spectrometer containing Turbo V electrospray ionization (ESI) source system united with Genius AB-3G nitrogen gas generator (Peak Scientific). The mass spectrometer was coupled with Agilent 1200 series HPLC system (Agilent Technologies) equipped with a degasser, a column oven, a binary pump and temperature-controlled auto sampler. The auto sampler was set at 4°C and column oven temperature was maintained in the range of 45 ± 1°C. The system control and data acquisition were executed by Analyst software (AB SCIEX, Applied Biosystems Inc., USA, version 1.5.1). Chromatographic separation was implemented by injecting 5 μL of samples on a Kinetex EVO C18 analytical column (100 × 2.1 mm, 100 Å, 5μm, Phenomenex Inc., CA, USA) under binary gradient conditions using mobile phase A (milliQ water containing 0.1% formic acid) and mobile phase B (acetonitrile containing 0.1% formic acid) with 400 μl/min flow rate. Analytes were eluted using the water-acetonitrile gradient i.e. 2% acetonitrile from 0-1 minute with linear increase to 85% from 1 to 4.8 minutes followed by linear increase to 99% till 5 minutes, hold at 99% acetonitrile up till 10.5 minutes.

The mass spectrometer was operated in multiple reaction monitoring (MRM) scan mode for identification of fragments in positive ionization mode at unit resolution. The optimized compound dependent instrument parameters for analytes with their MRM transitions are summarized in Figure 2 – supplement table 1. For all analytes, the ion spray voltage and source temperature/auxiliary gas temperature were set at 5000 V and 550°C respectively. Nebulizer gas (ion source gas 1, GS1) and auxiliary gas (ion source gas 2, GS2) were set at 30 psi. The curtain and collision-activated dissociation gases were set at 35 and 7 on an arbitrary scale. Data was processed using Multiquant software (AB SCIEX, USA, version 2.0), Microsoft Excel (Microsoft Inc., USA version 2013) and GraphPad Prism (LaJolla, CA, software version 8.4.2).

### Glycogen concentrations

Liver glycogen content was analysed using a Glycogen Assay Kit (Cayman Chemical, Michigan USA). Tissues were homogenised in assay buffer provided in the kit with the addition of 5 μL of protease inhibitor cocktail (Roche, Basel Switzerland) and centrifuged at 800 × *g* for 10 minutes at 4°C. The resulting supernatant was used for the glycogen assay, following the protocol outlined in the kit. The endpoint fluorescence assays were measured using a Flexstation 3 multimode plate reader (Molecular Devices, California USA). Glycogen concentrations were normalised to the protein concentration, measured using the PierceTM BCA Protein Assay kit (Thermo Fisher, Massachusetts USA).

### Immunofluorescence

The pancreas was dissected from SOD1_G93A_ and WT males at onset and mid-symptomatic stage of disease and incubated with 4% paraformaldehyde in 0.1 M phosphate buffer (pH 7.4, Sigma Aldrich, Missouri, USA) for 2 hours at room temperature, followed by 3 × 5 minutes washes in phosphate buffered saline (PBS, pH 7.4), and submersion in 15% and then 30% sucrose solution in PBS (pH 7.4). Pancreata were embedded in optimum cutting temperature (Sakura, California USA) and snap frozen in liquid nitrogen. Tissues were sectioned at 10 μm and dry mounted on Superfrost Plus slides (Menzel-Glaser, Braunschweig Germany) for quantification of α- and β-cell area.

Pancreas sections were rehydrated in PBS then blocked in PBS containing 5% bovine serum albumin (BSA) and 0.1% Triton X-100 for 1 hour at room temperature. Sections were incubated overnight at 4°C with guinea pig anti-insulin (1:50, Abcam, Massachusetts USA) and mouse IgG1 anti-glucagon (1:1000, Abcam) antibodies, followed by a 2-hour incubation at room temperature with Alexa Fluor 488 dye-conjugated goat anti-guinea pig (1:500, Invitrogen, Oregan USA) and Alexa Fluor 555 dye-conjugated goat anti-mouse IgG1 (1:1000, Invitrogen). All antibodies were diluted in PBS containing 1% BSA and 0.1% Triton X-100. Sections were then incubated at room temperature for 10 minutes with 4, 6-diamidino-2-phenylindole (DAPI, 1:25000, Thermo Fisher) diluted in PBS and mounted with Prolong Gold Antifade Medium (Invitrogen). Quantification of insulin and glucagon positive area was measured in 50 pancreatic islets for each sample. Staining procedures and image exposures were all standardized between genotypes and between sections. The mouse genotype was not made available to the researchers until the completion of the study.

### Real-time quantitative PCR

Total RNA was isolated from liver of WT and SOD1_G93A_ mice using RNeasy Lipid Tissue extraction kit according to manufacturer’s instructions (QIAGEN, CA, USA). The total RNA was purified from genomic DNA contamination then converted to cDNA using iScriptTM gDNA Clear cDNA synthesis kit according to manufacturer’s instructions (Bio-Rad, CA, USA). Commercially available SYBR® Green Glucose Metabolism Array consisting of 84 genes were used (Bio-Rad, CA, USA). Relative target gene expression to geometric mean of reference genes TATA-binding protein (Tbp), glyceraldehyde 3-phosphate dehydrogenase (Gapdh), and hypoxanthine-guanine phosphoribosyltransferase (Hprt) was determined using this formula: 2_−ΔCT_ where ΔCT = (Ct _(Target gene)_ – Ct _(Tbp, Gapdh and Hprt)_). Final measures are presented as relative levels of gene expression in SOD1_G93A_ mice compared with expression in WT mice. Pathway analysis was conduction using the “GENE2FUNC” function at FUMA GWAS web-based platform [41]. Gene lists examined included those showed changes in expression between WT and SOD1_G93A_ mouse liver samples (p < 0.1). The p-values were adjusted using Benjamini-Hochberg (FDR) multiple correction method. A pathway was considered significant at the adjusted p-value < 0.05 threshold.

### Statistical analysis

All analyses were performed using GaphPad Prism 8.4.2. All data generated from indirect calorimetry was analysed using a two-tailed student *t*-test with ages analysed separately. Blood glucose and hormone profiles over time for both ipGTT and ipITT were analysed using a two-way ANOVA followed by Bonferroni multiple comparison. For AUC, baseline hormone and glycogen concentrations and immunoreactive areas two-tailed student *t*-tests were used with different ages assessed separately. Two-tailed student *t*-test were also used for qPCR data, with all genes analysed individually. All correlations were assessed by Pearson correlation. All data is presented at mean ± SEM.

## Acknowledgements

The authors would like to sincerely thank technicians at the University of Queensland Biological Resources Animal Facilities for the animal care and husbandry. We also thank Maryam Shayegh for her technical support with genotyping the mice. JDL was supported by Motor Neuron Disease Research Australia (MNDRA) Postdoctoral Fellowship (PDF1604), and the research was funded by a grant from the National Health and Medical Research Council (NHMRC; Project grant APP1082271) and MNDRA (IG1930).

## Author’s contributions

TSM, TMW and JDL conceived the project. TSM and JDL designed the study. TSM performed the experiments with the assistance from VK and JNF. TMW and JDL provided supervisory and funding support for the project. All authors contributed to the analyses and interpretation of the data. TSM wrote the paper with contribution from JDL. All authors read and approved the final manuscript.

---

**Figure 2 – supplement figure 1.**
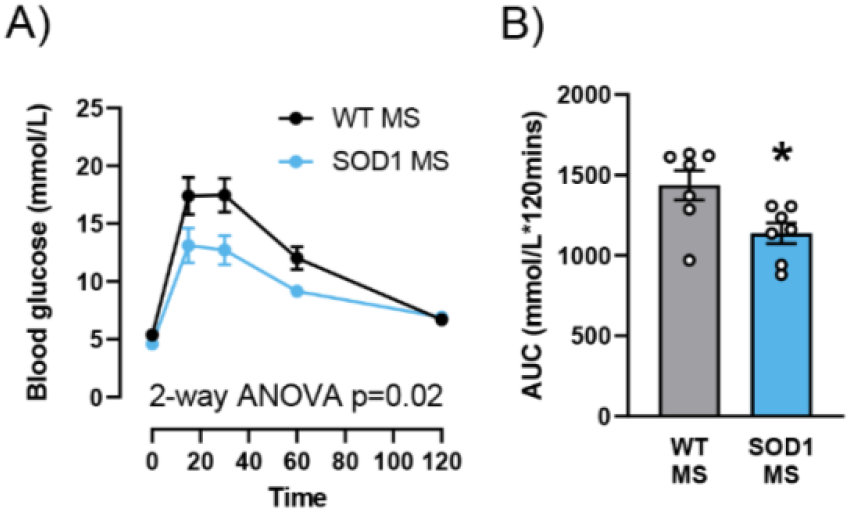
Female SOD1_G93A_ mice display a similar response to ipGTT in comparison to males. **(A)** Time course of blood glucose concentrations during a glucose tolerance test (ipGTT) following a 2 g/kg intraperitoneal injection of glucose at mid-symptomatic stage of disease in WT (black dots) and SOD1_G93A_ (blue dots) females. **(B)** The average area under the curve (AUC) calculated from the blood glucose time course for both WT (grey bars) and SOD1_G93A_ (blue bars) to measure degree of glucose tolerance. All data presented as mean ± SEM. *n* = 7, for all measurements. Two-way ANOVA results listed on time course graph is the overall significance between genotypes, Bonferroni *post-hoc* test was used to determine significant changes at specific time points. All bar graphs analysed by two-tailed student *t*-test. * p < 0.05.

**Figure 2 – supplement table 1.** Optimized MRM transitions and instrument parameters for quantification.

| Analyte | Q1 Mass (Da) | Q3 Mass (Da) | DP (eV) | EP (eV) | CEP (eV) | CE (eV) | CXP (eV) | RT (min) |
| --- | --- | --- | --- | --- | --- | --- | --- | --- |
| 2dOG | 270.2 | 91.1 | 41 | 6.5 | 16 | 31 | 4 | 5.7 |
|  |  | 124.1 |  |  |  | 25 |  |  |
|  |  | 55.1 |  |  |  | 77 |  |  |
| 2dOG6P | 350.3 | 91.1 | 41 | 6.5 | 16 | 31 | 4 | 5.8 |
|  |  | 124.1 |  |  |  | 25 |  |  |
|  |  | 65 |  |  |  | 77 |  |  |
| <sup>13</sup>C-glucose (Internal standard) | 292.07 | 91.1 | 36 | 9.5 | 24 | 35 | 4 | 6.4 |

**Figure 4 – supplement figure 1.**
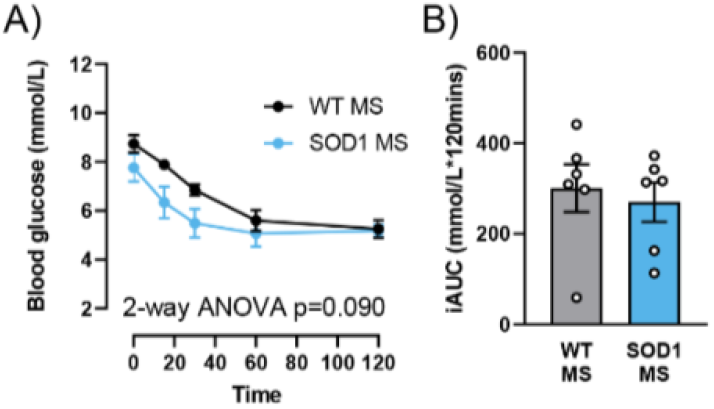
Female SOD1_G93A_ and WT littermates show a similar response to exogenous insulin. **(A)** Time course of blood glucose concentrations during an insulin tolerance test (ipITT) following a 0.5 IU/kg intraperitoneal injection of insulin at mid-symptomatic stage of disease in WT (black dots) and SOD1_G93A_ (blue dots). **(B)** The average inverse area under the curve (iAUC) calculated from the blood glucose time course for both WT (grey bars) and SOD1_G93A_ (blue bars). All data presented as mean ± SEM. *n* = 6, for all measurements. Two-way ANOVA results listed on time course graph is the overall significance between genotypes, Bonferroni *post-hoc* test was used to determine significant changes at specific time points. All bar graphs analysed by two-tailed student *t*-test.

**Figure 5 – supplement figure 1.**
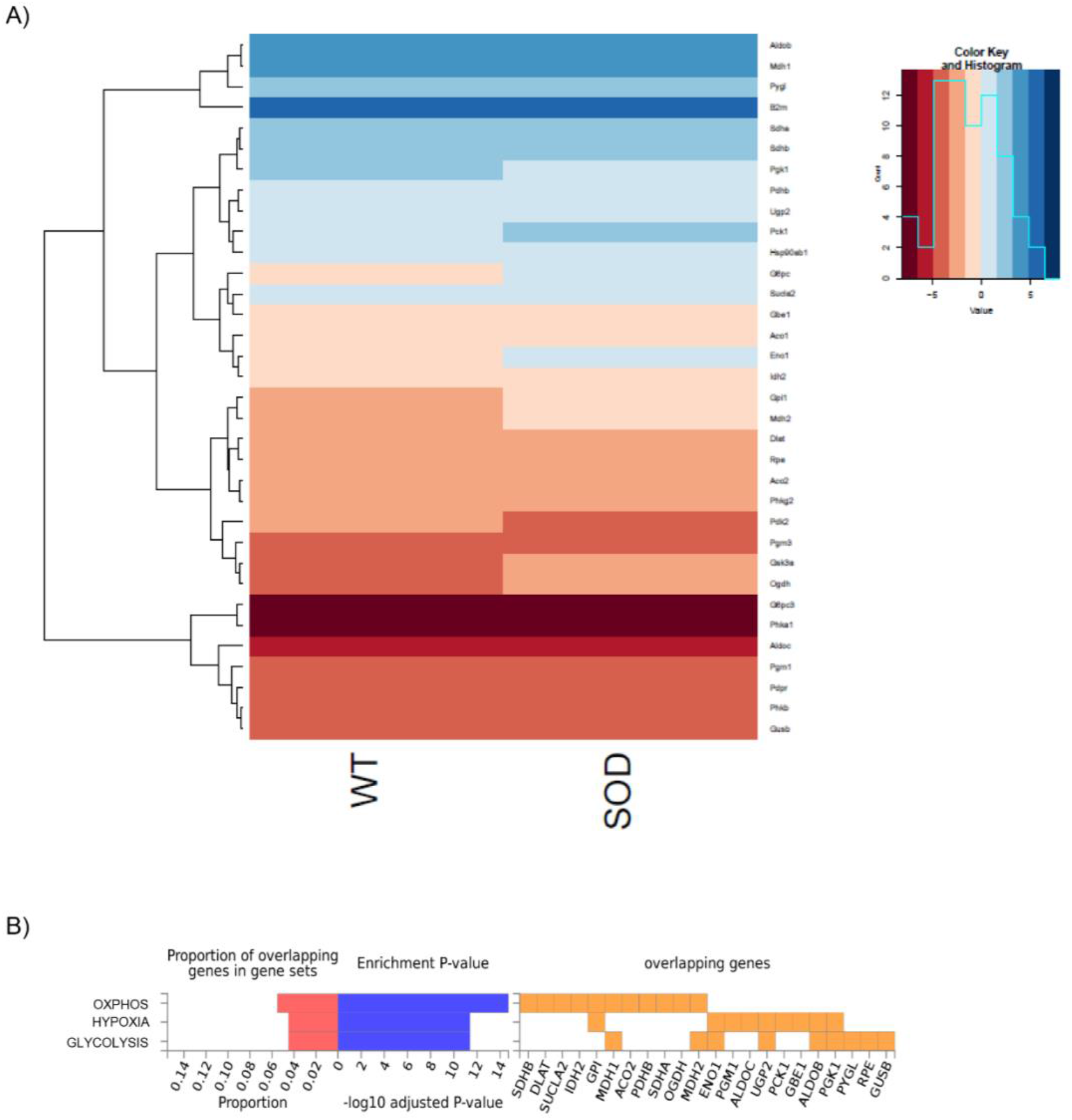
Three specific metabolic pathways are altered in liver of SOD1_G93A_ mice at mid-symptomatic stage. **(A)** Heatmap showing the gene expression profile of 34 genes with marked changes between WT and SOD1_G93A_ mouse liver samples (p < 0.1). The genes showing similar gene expression profiles are clustered. **(B)** Hallmark pathways enriched for genes with marked changes between WT and SOD1_G93A_ mouse liver samples (p < 0.1) using FUMA.

